# Computational Insights into the Allosteric Behavior of Mini Proinsulin Driven by C Peptide Mobility

**DOI:** 10.1101/2024.10.05.616825

**Authors:** Esra Ayan

## Abstract

**Background/aim:** The production of recombinant insulin remains challenging, particularly in enhancing refolding efficiency and bioactivity. Mini-proinsulin analogs, which involve reducing the length of the C-peptide, offer potential improvements in insulin production. This study aims to evaluate mini-proinsulin analogs’ design and receptor binding dynamics to optimize recombinant insulin production in *E. coli*.

**Materials and methods:** Mini-proinsulin analogs were engineered by replacing the 33-residue C-peptide with a pentapeptide sequence to improve refolding. The three-dimensional structure of mini-proinsulin was predicted using AlphaFold and performed docking analysis of mini-proinsulin analogs to the insulin receptor using AutoDock Tools, with comparisons made to previously available NMR-determined analog and the native insulin-insulin receptor complex. Normal Mode Analyses (GNM and ANM) were performed in detail to assess binding dynamics.

**Results:** *In silico* analyses revealed that mini-proinsulin analogs closely replicate the structural features of native insulin and display receptor binding dynamics similar to native insulin, though they follow distinct receptor interaction paths.

**Conclusion:** All analysis suggests that C-peptide mobility may contribute to the allosteric behavior observed in mini-proinsulin analogs during receptor interaction.

## 1. Introduction

Insulin, a polypeptide hormone secreted by pancreatic β-cells, comprises A and B chains. The chains are connected by two interchain and an intrachain disulfide bond [1]. The hormone is initially produced as a single-chain precursor, preproinsulin, which undergoes proteolytic cleavage in the pancreas to forge its active form. In the human body, preproinsulin has signal peptide, B-peptide, C-peptide and A-peptide. However, in the recombinant production, preproinsulin has fusion tag, B-peptide, C-peptide and A-peptide. In the absence of signal peptide in human body or without fusion tag in recombinant production, insulin is termed as proinsulin [1].

*Escherichia coli* (*E. coli*) is typically employed in the production of recombinant human insulin, where preproinsulin is synthesized in the bacterial cytoplasm and accumulates as inclusion bodies within a fusion protein. Following the CNBr administration to the fusion product and sulfonation of the released proinsulin, the refolding process of S-sulfonated proinsulin can proceed with relatively high efficiency [2]. After tryptic digestion of proinsulin, mature insulin, which consists of B-peptide and A-peptide alone, is produced, releasing the C-peptide, whose length varies between 26 and 38 amino acid residues across different animal species [3]. Mature insulin is vital for regulating blood glucose levels, while the C-peptide, co-secreted with insulin, exhibits biological activity, including its ability to repair the diminished Na^+^-K^+^ dependent ATPase activity observed in diabetic conditions [4, 5].

Proinsulin displays only 2% of the receptor binding capacity compared to mature insulin [6]. Conversely, an insulin analog that was directly cross-linked between the C-terminus of the B chain and the N-terminus of the A chain exhibited a near-total loss of bioactivity despite its crystal structure being nearly indistinct from that of native insulin [7, 8]. It indicates that the length of the cross-link is critical in determining insulin’s bioactivity. Still, a study conducted by Brems et al. (1991) systematically investigated the effect of cross-linking between the B and A chains, revealing that a specific cross-link length is critical for maintaining bioactivity [9]. Dual-chain analogs, such as conventional insulins, present significant challenges due to their susceptibility to fibrillation. Glidden et al. developed a formulation of a specific cross-link length mini-proinsulin analog that exhibits enhanced resistance to self-assembly at high concentrations. The conclusions reveal that the residues responsible for receptor binding are localized on both the B and A chains. Insulin has activity only when it possesses sufficient flexible C-peptide to adopt an optimal orientation, allowing the interacting residues to align correctly with the receptor.

The three-dimensional structure of insulin indicates that a peptide linker could connect the A and B chains significantly shorter than the 33-residue C-peptide [10, 11]. However, it remains novel whether reducing the length of the C-peptide would enable the resulting mini-proinsulin to exhibit receptor interaction dynamics comparable to those of mature insulin. In related research, mini proinsulin was engineered to enhance the refolding efficacy of the insulin precursor and was also facilitated recombinant production by increasing productivity in *E. coli* [6]. These parameters are still challenging for recombinant insulin production in industrial applications. The findings revealed a 20–40% improvement of mini proinsulin use in refolding efficiency compared to standard proinsulin. The mini proinsulin exhibited substantial receptor binding activity, at least 50% compared to native insulin [6, 11]. Interestingly, the recent study shows that unlike conventional insulin molecules, which are prone to thermal degradation, aggregation, and stability issues, the single-chain mini-proinsulin analog demonstrates greater stability and production efficiency [10]. Specifically, the mini-proinsulin that incorporates a flexible linker enhances refolding efficiency, thereby supporting cost-effective manufacturing. Moreover, the activity of the mini-proinsulin that closely resemble native insulin underscores their potential to deliver comparable biological activity, strengthening their therapeutic prospects.

Here, the 33-residue C-peptide is replaced with a pentapeptide sequence designed to form a short turn, thereby promoting refolding. The novel designer mini proinsulin, R22B and P28B, are substituted by K22B and D28B, respectively, to minimize aggregate formation *in solution*. Additionally, the substitution of N60A for G60A is performed to enhance stability in acidic conditions. The three-dimensional structure of mini proinsulin predicted using AlphaFold (AF), has been perform of docking with the insulin receptor. Similarly, the mini proinsulin, whose structure had been previously determined through an NMR study in the literature (PDB ID: 1EFE), has been also performed of docking with the receptor. Both docking structures are compared with the experimental insulin-insulin receptor complex (PDB ID: 6VEP) [12] using *in-silico* analyses. Previously available experimental results had demonstrated a high ratio of insulin moiety in the fusion protein as well as enhanced the refolding capacity of mini proinsulin [6, 10]. The *in-silico* analysis suggests that despite mini proinsulin analogs having nearly identical intrinsic dynamics post-binding mechanism to that of native insulin, they display allosteric behavior in receptor binding due to probably C-peptide mobility. Collectively, further well-optimized mini-proinsulin analogs might be used as final products to produce recombinant insulin in E. coli.

## 2. Materials and methods

### 2.1. Protein Structures

The AlphaFold prediction of novel designer mini-proinsulin structure (FVNQHLCGSHLVEALYLVCGEKGFFYTDKTRRYPGDVKRGIVEQCCTSICSLY QLENYCG) has been used for this study along with the mini-proinsulin structure from the Protein Data Bank (PDB) under the 1EFE accession code. Additionally, insulin-insulin receptor structure from PDB (ID: 6VEP) has been used for this study to perform docking analyses between mini-proinsulin and insulin receptor as well as to compare receptor interaction dynamics between genuine insulin-insulin receptor and mini-proinsulin-insulin receptor. An unbound native insulin structure has been used to compare dynamicity in isolation between native insulin from PDB (ID: 3I40) and mini-proinsulin analogs.

### 2.2. Structure Prediction

The novel designer mini-proinsulin has been calculated as a monomer using AlphaFold (AF) with MMseqs2 v1.5.5. The structure prediction process followed the standard approach outlined in the AlphaFold paper [13], which involves five key steps: multiple sequence alignment (MSA) construction, template search, inference with five models, ranking of models based on the mean predicted local distance difference test (pLDDT), and constrained relaxation of the predicted structures. For the novel mini-proinsulin, AF has been executed with the model preset set to “monomer” using the following command:

~~~
alphafold --fasta_paths=<FASTA file path> --model_preset=<MONOMER> --
output_dir=<OUTPUT directory> --db_preset=full_dbs --use_gpu_relax --
max_template_date=<MAX template date>
~~~

The structure with the highest prediction accuracy has been selected for further analysis. Molecular images have been generated by PyMol version 2.3.0 (Schrödinger, LLC, New York, NY, USA).

### 2.3. Docking analysis

AutoDock and AutoDock Tools [14] were used to optimize docking parameters in precision. The necessary coordinate files and related information were converted from PDB format to PDBQT format, which includes atom and bond types, partial atomic charges, and other relevant data. After loading the insulin receptor structure, the target file was prepared by adding hydrogen atoms, correcting bond orders, and renumbering residues. Kollman charges, derived from quantum mechanical calculations, were applied to assign appropriate partial charges to the receptor. Similarly, the mini-proinsulin ligand’s coordinate files were processed in PDBQT format, with hydrogen atoms added, bond orders corrected, and residues renumbered. Gasteiger charges were applied to the ligand to facilitate the docking procedure. A grid parameter file was prepared to calculate grid maps for interaction energies across various atom types, essential for calibrating the docking procedure. A grid box was generated to define the binding site coordinates for the search engine. A docking parameter file was created to execute the docking using AutoDock Tools. The following commands were used to run AutoGrid and AutoDock:

~~~
autogrid4.exe -p dock.gpf -l dock.glg &
autodock4.exe -p dock.dpf -l dock.dlg &
~~~

The resulting dlg file contained detailed information about the docking simulations, including the calculated binding energy in kcal/mol. The best conformations had binding energies of −6.35 kcal/mol for the nM2PI-IR co-complex and −4.46 kcal/mol for the M2PI-IR co-complex. Additionally, the root-mean-square deviations (RMSD) of atomic positions of nM2PI-IR and M2PI-IR co-complexes from the reference co-complex (PDB ID: 6VEP) were 0.329 Å and 0.312 Å, respectively.

### 2.4. Gaussian Network Model (GNM)

The Gaussian Network Model (GNM) analysis was performed using ProDy [15]. GNM analysis was completed on both mini-proinsulin variants and native insulin in isolation, as well as on docked co-complexes (nM2PI-IR and M2PI-IR) and the native insulin receptor-insulin (IR-INS) co-complexes. For all structures, Cα atoms were operated to construct contact maps. The cutoff distance was set to 8.00 Å, and N − 1 nonzero normal modes were obtained, where N represents the number of Cα atoms. Cross-correlations between residue fluctuations were calculated based on the cumulative 10 slowest GNM modes. The squared fluctuations for each structure were calculated across the weighted 10 fastest modes to compare their local motions and dynamic behavior. Graphics illustrating the normalized results were generated using Matplotlib.

### 2.5. Anisotropic Network Model (ANM)

ANM (Anisotropic Network Model) analysis was conducted on the docking structures of nM2PI-IR, M2PI-IR, and the native INS-IR co-complex (PDB ID: 6VEP). A cutoff distance of 12.0 Å and the default spring constant of 1.0 were used. The ANM modes were focused on individual mode 5. The resulting protein motions were visualized using the NMWiz tool within the VMD software.

## 3. Results

### 3.1. Novel mini-proinsulin mimics the structural landscape of native insulin

The novel mini-proinsulin (hereafter referred to as nMPI2) was designed by reducing the C-peptide from 33 residues to a 5-residue mini-C peptide, along with substituting Arginine with Lysine at position 22 of Chain B (hereafter referred to as R22B, K22B) and Proline with Aspartic acid at position 28 (hereafter referred to as P28B, D28B). Additionally, Asparagine at position 60 of Chain A was replaced by Glycine (hereafter referred to as N60A, G60A) (Fig. S1). To obtain the three-dimensional structure of nMPI2 (Fig. 1A), an AF prediction was conducted, as described in the Methods section. The experimentally determined mini-proinsulin structure (hereafter referred to as MPI2) (PDB ID: 1EFE) (Fig. 1B), resolved by NMR techniques, was used to compare with the predicted nMPI2 structure, indicating a perfect superposition with a root-mean-square deviation (RMSD) of up to 1.2 Å. Furthermore, the structure of human insulin (hereafter referred to as INS) (PDB ID: 3I40), determined through X-ray crystallography, was used to compare the dynamic deviations between the mini-proinsulin and native INS in isolation. A highly consistent RMSD matching up to 0.4 Å was observed between the mini-proinsulin analogs and native INS (Fig. 1C).

**Figure 1.**
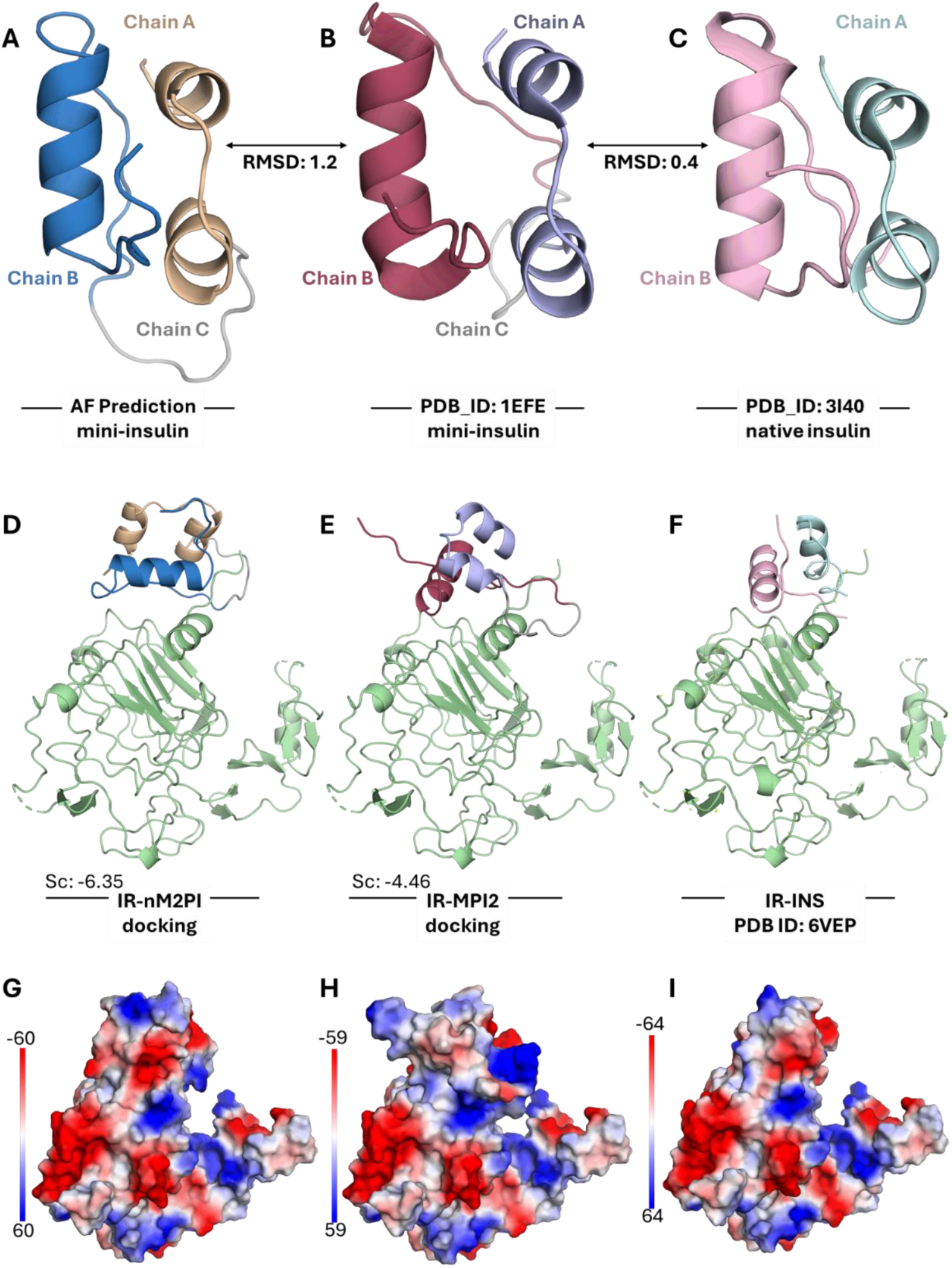
Structural representation of mini-analogs for calculated and experimental co-complexes. (A) AlphaFold prediction of the novel designer mini-proinsulin (nMPI2). (B) Experimental structure of mini-proinsulin from the Protein Data Bank (PDB ID: 1EFE), referred to as MPI2. (C) Experimental structure of native insulin from the PDB (ID: 3I40). RMSD values between the structures are indicated with bidirectional arrows. (D) Docking co-complex of the experimental insulin receptor (IR, shown in green) and nMPI2, indicating a relatively higher affinity score. (E) Docking co-complex of the experimental IR (green) and MPI2, indicating a relatively lower affinity score. (F) Experimental IR-INS Cryo-EM co-complex from the PDB (ID: 6VEP). (G, H, I) Poisson-Boltzmann electrostatics of the docking and experimental co-complexes, illustrating the charge-smoothed protein contact potential. Red regions represent areas of negative electrostatic potential, blue regions indicate areas of positive electrostatic potential, and white (or neutral) regions represent areas with near-neutral electrostatic potential.

Docking analysis, as described in the Methods section, was conducted to evaluate if the mini-proinsulin analogs could effectively bind to the insulin receptor. This step was essential to identify potential binding modes and to serve as a foundation for further dynamic analyses since the dynamics between mini-proinsulin and insulin receptor are still unknown in the literature. This analysis included a comparison with the experimental structure of the insulin-insulin receptor co-complex (hereafter referred to as IR-INS) (PDB ID: 6VEP) (Fig. 1D-F). According to the molecular docking results, nMPI2 exhibited a relatively higher affinity (−6.35) for the αCT domain of the insulin receptor (IR), whereas MPI2 showed a lower affinity (−4.46). Both mini-proinsulin analogs demonstrated ideal alignment with the experimental IR-INS structure, with RMSD of 0.319 Å for the IR-nMPI2 co-complex and 0.312 Å for the IR-MPI2 co-complex.

Electrostatic forces can give required information about protein-protein interactions and protein stability [16]. The insulin-binding region for each co-complex was surrounded by an acidic and neutral region concentrated with stable negatively charged and uncharged residues in the presence even of the charge substitutions in the nM2PI conformer. The electrostatic distribution here highlights areas where charged interactions, such as hydrogen bonds or ionic interactions, might occur between the nM2PI and the IR (Fig. 1G). The charge distribution might be crucial for receptor binding and overall affinity. However, the electrostatic distribution, while similar, may show slight differences in charge regions compared to the IR-nM2PI co-complex, reflecting lower affinity or less optimized electrostatic interaction between IR and M2PI co-complex (Fig. 1H). Notably, certain blue and red areas seem less extensive or less aligned with the receptor, potentially explaining the lower binding affinity, which is highly compatible with the docking score of the IR-M2PI co-complex (Fig. 1E). The electrostatic potential map of the experimental IR-INS co-complex demonstrates their natural, highly optimized charge interaction (Fig 1I). The alignment and balance of positive and negative regions are more uniform and organized, likely indicating stronger, more stable interactions than the docked mini-proinsulin analogs. Still, Panel G in Figure 1 illustrates that the IR-nM2PI co-complex attempts to mimic the natural electrostatic landscape of native insulin (Fig. 1I) but with varying degrees of success. It is essential that there are highly similar electrostatic charge distributions between IR-nM2PI and experimental co-complexes (Fig G and Fig. I), whereas potential surface charge redistributions may force shape perturbations of a protein by changing hydrogen bonds and salt bridges.

Additionally, an overlay analysis of mini-proinsulin forms with native insulin was performed, and residue-by-residue differences were examined (Fig. 2). The results indicate that nM2PI exhibits allosteric behavior compared to the native form. This analysis provides a comprehensive view of the primary binding mode of site 1 and site 2 on the insulin receptor (Site 1: Gly A1, Ile A2, Val A3, Glu A4, Tyr A19, Asn A21, Gly B8, Ser B9, Leu B11, Val B12, Tyr B16, Phe B24, Phe B25, Tyr B26; Site 2: Thr A8, Ile A10, Ser A12, Leu A13, Glu A17, His B10, Glu B13, Leu B17) [17], highlighting its interactions with the L1 domain (Asp 12, Arg 14, Asn 15, Leu 37, Phe 39, Lys 40, Phe 64, Glu 97, and Lys 121) and the αCT domain (Asp 707, His 710, Phe 714, Pro 716, Arg 717, and Pro 718) [12, 18]. Notably, nM2PI establishes novel interaction networks with the insulin receptor, particularly through its Chain B (Fig. 2K).

**Figure 2.**
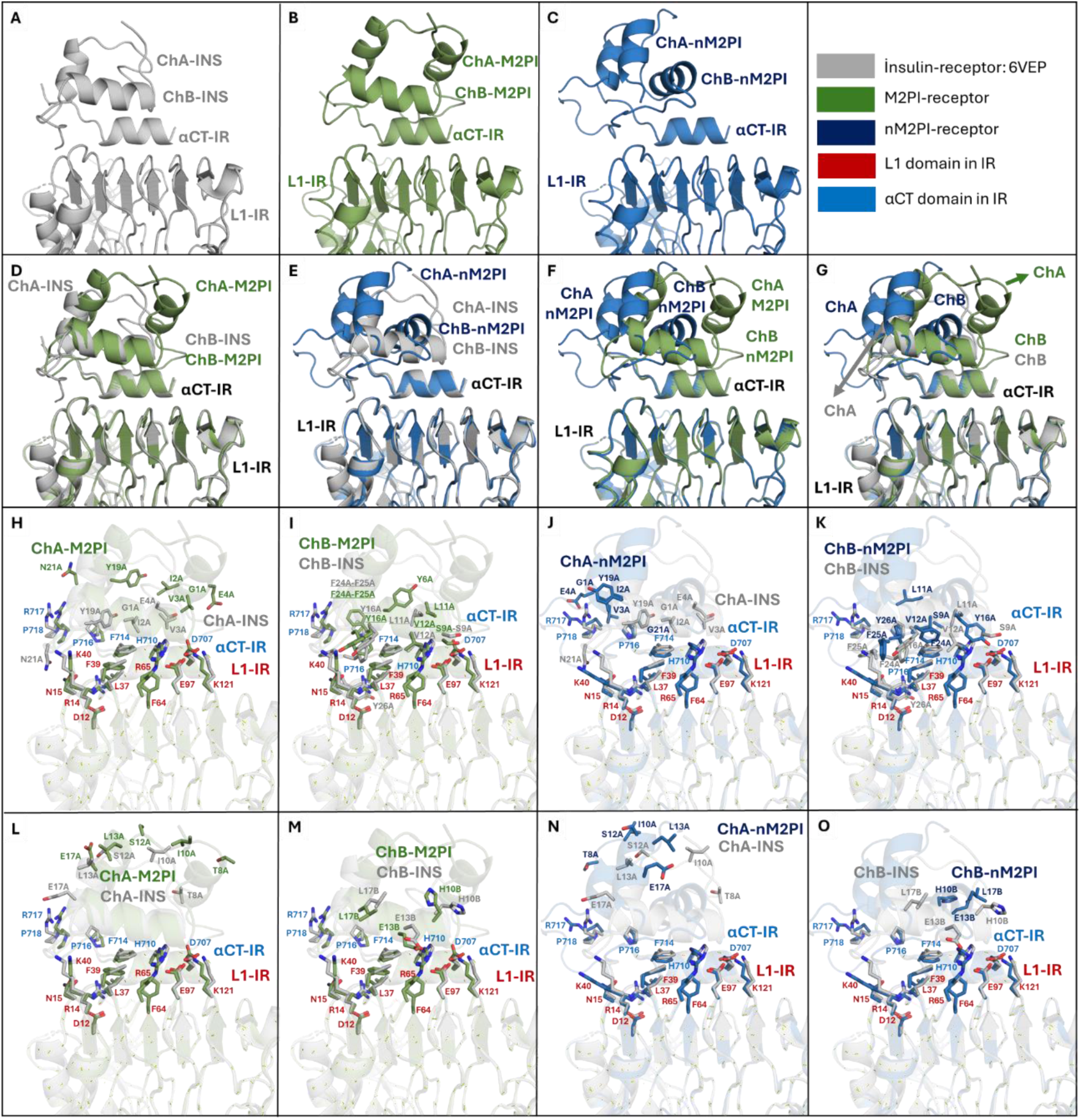
Structural overlays aligning the mini-proinsulin analogs with the insulin receptor to clarify probable binding mode differs. Panel A illustrates the native insulin-receptor (INS-IR) binding mode, obtained from the PDB structure (ID: 6VEP). Panel B shows a docking co-complex between the insulin receptor (extracted from the 6VEP structure) and mini-proinsulin (M2PI-IR) derived from the 1EFE PDB structure. This complex shows an alternate binding conformation, likely due to the presence of the C-peptide. Panel C presents another docking co-complex between the insulin receptor (6VEP) and a novel designer mini-proinsulin (nM2PI-IR) modeled using the AlphaFold tool, which demonstrates significant conformational changes in receptor binding. Panel D compares the superposition of INS-IR and M2PI-IR co-complexes, revealing slight differences in the binding of Chain A (ChA). Panel E shows a superposition of INS-IR and nM2PI-IR, highlighting substantial differences in both Chain A (ChA) and Chain B (ChB) conformations. Panel F compares the M2PI-IR and nM2PI-IR co-complexes, demonstrating pronounced differences in the ChA and ChB peptides. Panel G overlays INS-IR, M2PI-IR, and nM2PI-IR co-complexes, showing that both chains of nM2PI-IR exhibit significant distinctions compared to the other co-complexes. Panel H focuses on Chain A (IR binding site 1: G1A, I2A, V3A, E4A, Y19A, and N21A) of the M2PI analog, which exhibits lower binding affinity than native insulin. Panel I highlights that Chain B (IR binding site 1: GB8, SB9, LB11, VB12, YB16, FB24, FB25, YB26) of M2PI demonstrates strong binding affinity to the insulin receptor, comparable to native insulin’s Chain B. Panel J indicates that residues G1A, I2A, V3A, E4A, and Y19A of Chain A in nM2PI are localized on the left side of the co-complex relative to native insulin, resulting in a strong affinity for residues P716, R717, and P718 in the aCT domain of the insulin receptor. Panel K shows that, except for residue L11B, the other IR-binding residues of Chain B in nM2PI exhibit strong affinity to both the αCT and L1 domains of the insulin receptor, albeit with distinct binding patterns compared to native insulin. Panel L and N focuses on Chain A (IR binding site 2: T8A, I10A, S12A, E13A, and L17A) of the M2PI and nM2PI analogs, respectively, which those residues in both forms exhibit similar localization on IR compared to native insulin. Panel M and O highlights that Chain B (IR binding site 2: H10B, E13B, and L17B) of M2PI and nM2PI forms demonstrate those residues of both forms shows close proximity to αCT domain of IR.

### 3.2. Mini proinsulins exhibit enhanced stability compared to the unbound state

Protein binding is determined by setting novel noncovalent interactions between specific residues of two proteins. These interactions can impact residues located at distant sites by disrupting current communication paths and creating new ones. The reorganization has the potential to significantly change the conventional residue communication network [19]. Such modifications can be effectively identified and analyzed using the Normal Mode Analysis (NMA). GNM and ANM, rooted in NMA, effectively predict large-scale protein movements in slow modes and localized dynamics in fast modes. Regions with higher fluctuations in slow modes indicate mobile segments, whereas those with reduced fluctuations represent hinge points essential for protein functionality. In fast modes, residues exhibiting peak fluctuations often signify structurally constrained hot spots that play critical roles in binding or folding. Additionally, examining variations in residue dynamics across all modes offers valuable insights into the functional mechanisms and allosteric regulation of proteins [20]. Here we performed cross-correlations between residue fluctuations of weighted 10 slowest GNM modes for each case (IR-nM2PI; IR-M2PI; IR-INS) (Fig. 3). Cross-correlation for calculated IR-nM2PI co-complex indicated nM2PI almost behaves as a rigid block while binding to IR, all residues practically moving along the same mode axis (cross-correlations close to 1 (red)) (Fig. 3A-B), which is highly compatible with diminished B-factor (Fig. 3C) of sliced nM2PI (Fig. 3B) compared to greater mobile B-factor (Fig. 3E) of nM2PI in isolation (Fig. 3D). Namely, nM2PI in isolation reveal greater variations in the movement directions of various residues or groups of residues, with some moving in opposite directions and being separated by hinge regions (cross-correlations close to 1 (blue)) (Fig. 3D). Cross-correlation for calculated IR-M2PI co-complex also shows the similar characteristic with the calculated IR-nM2PI co-complex (Fig. 3F-J). However, experimental IR-INS co-complex indicated more optimized and strongest correlation compared to Panel A and Panel F in Figure 3 (Fig. 3K-L). Additionally, native insulin exhibits the lowest mobility once binding to the IR, suggesting that the interaction is characterized by maximal affinity and structural stability (Fig. 3M) compared to unbound conformer (Fig. 3N and Fig. 3O). Computational analyses were also performed using a novel mini-proinsulin analog, (hereafter referred to as SCI), which features a 6-amino acid linker instead of the pentapeptide (Fig. S2) [10]. The crystal structure of SCI was superposed with the nM2PI structure, yielding a RMSD of 0.91 Å (Fig. S2A). Cross-correlation analysis revealed a higher level of collectivity and cooperativity in SCI during binding to the IR (Fig. S2B-C), while hinge-like behavior was observed, dissecting the dynamic domains of SCI when analyzed in isolation (Fig. S2D). These findings are consistent with the intrinsic dynamics observed in the native IR-INS co-complex (Fig. 3K,L,N).

**Figure 3.**
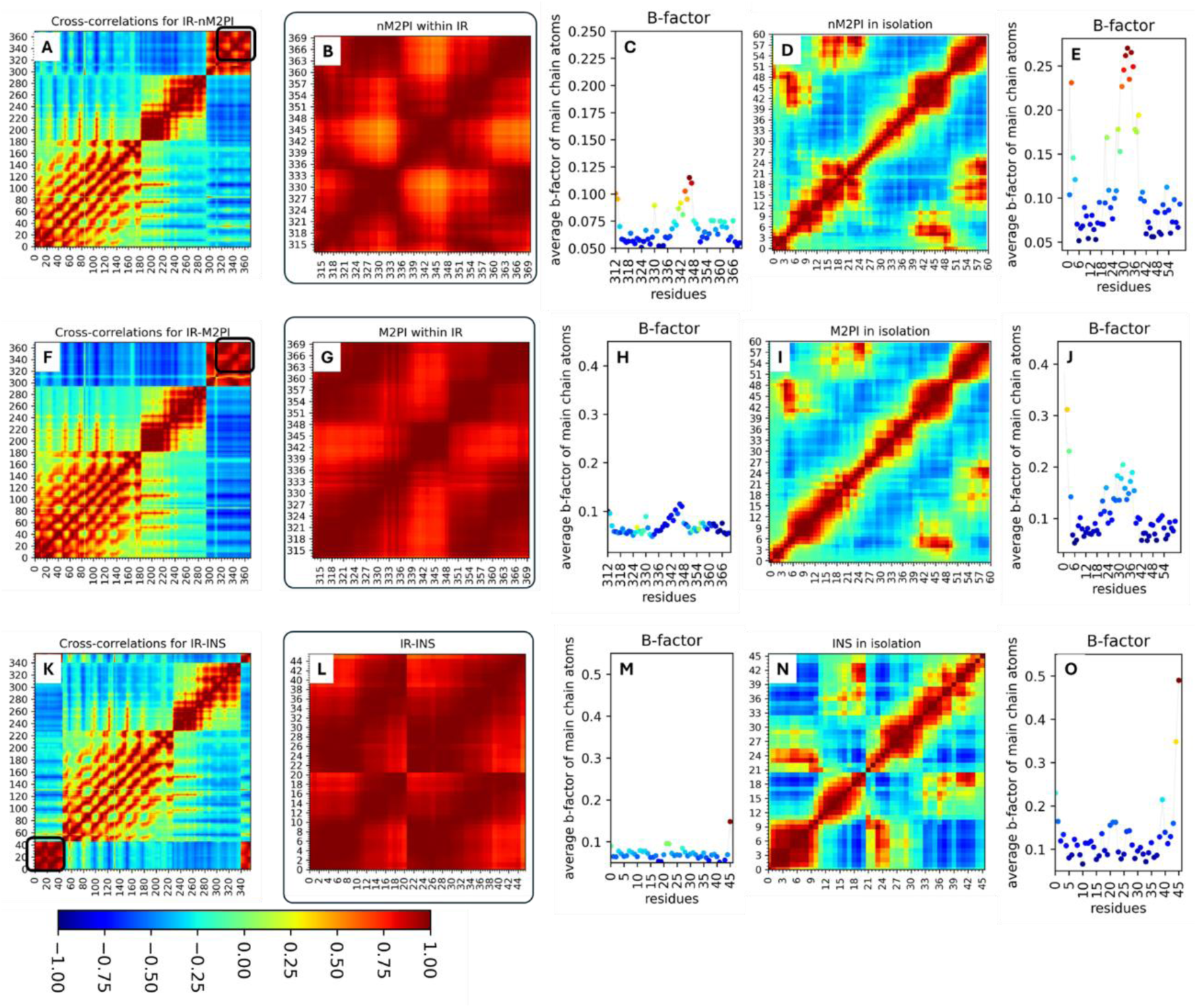
Cross-correlation and calculated B-factor by GNM comparisons between calculated and experimental co-complexes. (A) Cross-correlation for IR-nM2PI. Highlight square indicated docking insulin monomer’ cross-correlation. Residue pairs exhibiting correlated motion, moving in the same direction, are depicted in red regions (C_ijorient_ > 0), while those exhibiting anticorrelated motion, moving in opposite directions, are represented in blue regions (C_ijorient_ < 0). Uncorrelated residue pairs are shown in green (C_ijorient_ = 0.0), as indicated by the color bar on lower left corner. (B) Cross-correlation for nM2PI monomer within IR-nM2PI co-complex. (C) B-factor of nM2PI monomer within IR-nM2PI co-complex. (D) Cross-correlation for nM2PI monomer in isolation. (E) B-factor of nM2PI monomer in isolation. (F) Cross-correlation for IR-M2PI. (G) Cross-correlation for M2PI monomer within IR-M2PI co-complex. (H) B-factor of M2PI monomer within IR-M2PI co-complex. (I) Cross-correlation for M2PI monomer in isolation. (J) B-factor of M2PI monomer in isolation. (K) Cross-correlation for IR-INS. (L) Cross-correlation for INS monomer within IR-INS co-complex. (M) B-factor of INS monomer within IR-INS co-complex. (N) Cross-correlation for INS monomer in isolation. (O) B-factor of INS monomer in isolation.

Collectively, cross-correlation analyses show that native insulin provides the most robust and conformationally stable form while binding to IR (Fig. 3L-M). While nM2PI, M2PI and SCI exhibit comparable correlation profiles, native insulin presents a more refined optimization of structural integrity. These observations are verified by *B*-factor analyses, where native insulin exhibits the lowest dynamic mobility post-receptor binding (Fig. 3M). Although the novel and alternative mini-proinsulin analogs have relatively higher residue mobility after receptor docking, they display markedly improved structural rigidity compared to their unbound conformer (Fig. 3B,D,G,I). Thus, native insulin displays the highest binding affinity and structural fidelity, while the mini-proinsulin analogs, although demonstrating a similar binding paradigm, do not perform the same level of stabilization and optimization, confirming mini-proinsulin demonstrates receptor binding efficacy similar to native insulin [6, 10].

### 3.3. ANM modes support nM2PI shows post-binding stability

The ANM analysis can illustrate that intrinsic residue fluctuations in C-peptide mobility drive the allosteric behavior of mini-proinsulin variants post-binding to IR. ANM provides insight into residue motion directions over 3D structure by calculating theoretical displacements of residues. Especially, the absence of a full-length C-peptide represents a significant structural modification. Therefore, ANM was employed in this study to investigate the critical allosteric effects in nMP12 and MP12, specifically focusing on Mode 5. This approach allowed us to characterize, in detail, the interactions between local residue fluctuations and global motion patterns across the three-dimensional structure, as captured through simulation vectors. The direction of motions, derived from the eigenvectors of Mode 5, was mapped onto the IR-INS, IR-M2PI, and IR-nM2PI co-complexes, with a color gradient representing the amplitude of the residue fluctuations. In the native IR-INS interaction, the correlation map reveals a well-coordinated residue motion pattern with distinct regions of coherent dynamics (Fig. 4A). The movements suggest that INS displays highly optimized collective motions with the receptor, indicative of a well-tuned allosteric communication network captured by the ANM analysis. Instead, the IR-M2PI appears more disordered and less coordinated than the INS conformer (Fig. 4C). Yet, the correlation map of IR-nM2PI unveils relatively concerted and robust correlations than M2PI (Fig. 4E). The red and blue regions are more coherent, indicating relatively coordinated collective motions and a well-optimized dynamic interaction with the receptor, suggesting nM2PI favors a dynamic modulation closely similar to INS. The INS direction of motion within the IR-INS co-complex are more dominant in Mode 5 (Fig. 4B) compared to the IR-M2PI and IR-nM2PI co-complexes (Fig. 4D-F). Notably, Chain B in the M2PI and nM2PI analogs exhibits increased mobility, indicating less restricted dynamics in the region. Still, M2PI exhibits relatively weaker collective dynamics and more irregular motion patterns (Fig. 4D), suggesting reduced post-binding stability compared to INS conformer.

**Figure 4.**
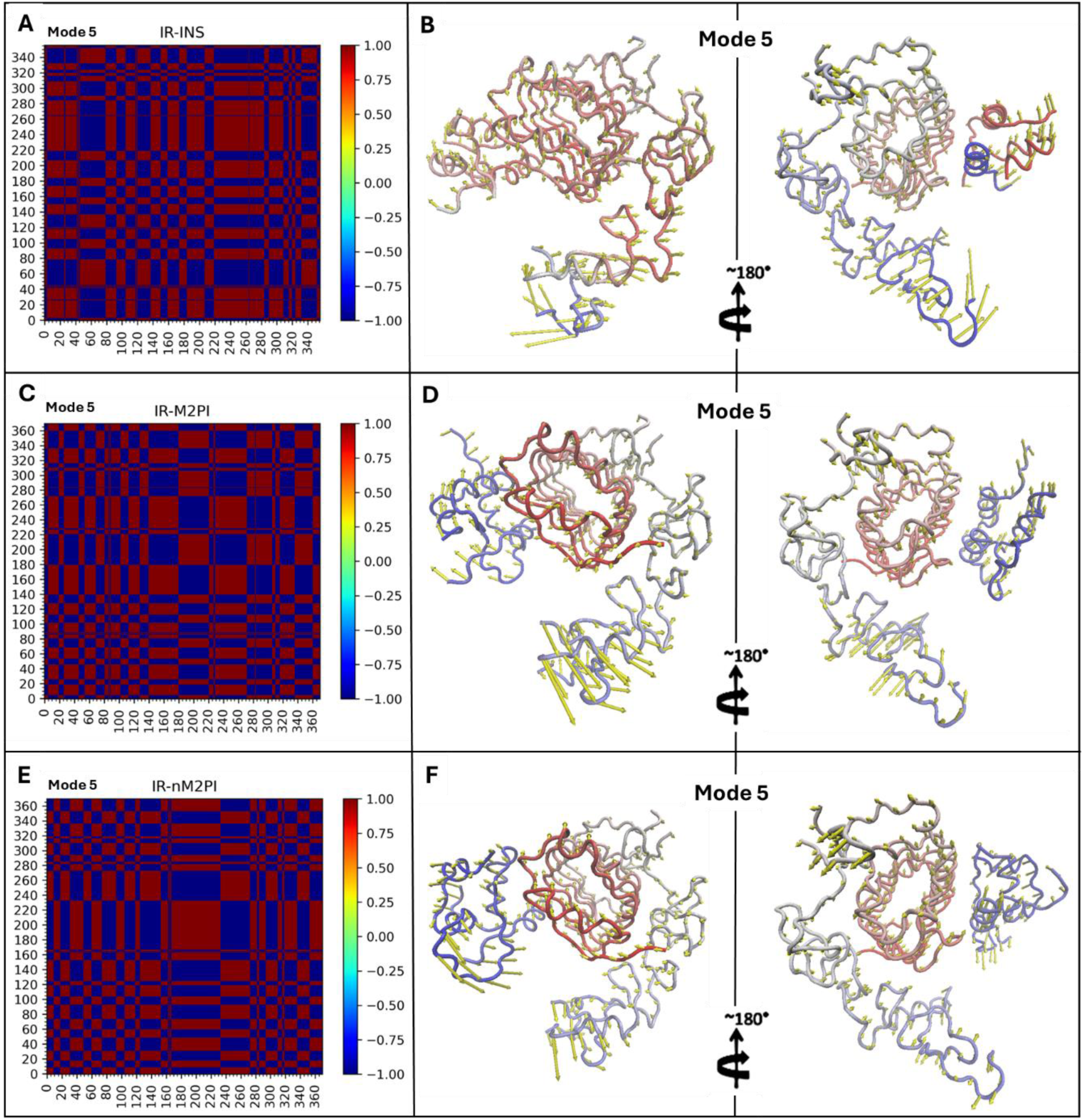
Comparison of individual 5 slow ANM mode direction of motions in INS, M2PI, and nM2PI bound to the IR. (A) Cross-correlations of IR-INS map by ANM. Residue pairs exhibiting correlated motion, moving in the same direction, are depicted in red regions (C_ijorient_ > 0), while those exhibiting anticorrelated motion, moving in opposite directions, are represented in blue regions (C_ijorient_ < 0) (B) Mode 5 of IR-INS is drawn on the structure. Ribbon representation was rendered blue to gray and red coloring, representing high to low mobility (C) Cross-correlations of IR-M2PI map by ANM. (D) Mode 5 of IR-M2PI is drawn on the structure. (E) Cross-correlations of IR-nM2PI map by ANM. (F) Mode 5 of IR-nM2PI is drawn on the structure.

### 3.4. Local motions reveal unprecedented hotspot residues in mini proinsulins

GNM yield precise estimations of localized dynamics within the fast vibrational modes. Residues exhibiting peak fluctuations in the fast modes are potential hot loci and are structurally constrained and play pivotal roles in folding and ligand binding processes. Variations in residue motion profiles offer valuable insights into the protein’s functional mechanisms and allosteric modulation. To assess and compare local dynamics, squared fluctuations across the top 10 weighted fastest GNM modes were computed for both predicted co-complexes compared experimental IR-INS interactions (Fig. 5). Additionally, to assess collective motion between only αCT domain of IR, which is critical part of the insulin binding, and insulin variants (M2PI, nM2PI, and INS), performed cross-correlation over the 10 slowest GNM modes.

**Figure 5.**
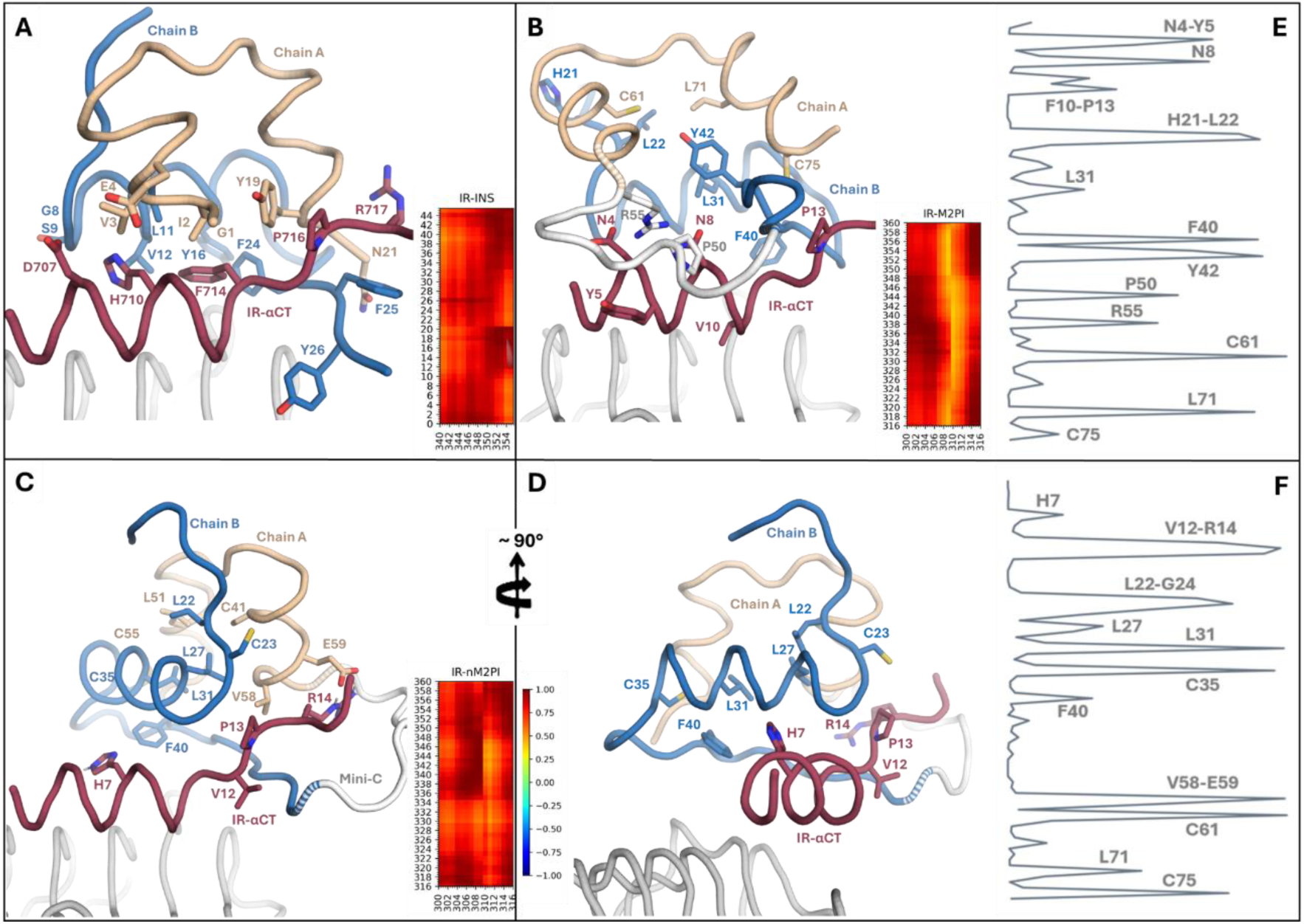
Structural localization of potential receptor interaction residues and their GNM analysis. (A) Native αCT-INS interaction residues adapted from PDB structure under the 6VEP accession code. Also, cross-correlation analysis between αCT domain and INS to emphasize their strong collective motion. (B) αCT-M2PI interaction residues determined by 10 fastest GNM modes and cross-correlation analysis between αCT domain and M2PI to emphasize their relatively strong collective motion. (C) αCT-nM2PI interaction residues determined by 10 fastest GNM modes and cross-correlation analysis between αCT domain and nM2PI to emphasize their strong collective motion. (D) Turn panel C to 90 degrees for inspecting the detailed interaction between the αCT-nM2PI complex. Residue pairs exhibiting correlated motion, moving in the same direction, are depicted in red regions (C_ijorient_ > 0), while those exhibiting anticorrelated motion, moving in opposite directions, are represented in blue regions (C_ijorient_ < 0). Uncorrelated residue pairs are shown in green (C_ijorient_ = 0.0), as indicated by the color bar in those panels. (E, F) Ten fastest GNM modes of αCT-mini analogs interactions. Potential hot loci have been labelled with relevant residues.

Compared to the experimental IR-INS interaction (Fig. 5A), the local dynamics between the αCT domain of the IR and the mini-proinsulin analogs (Fig. 5B-F) reveal distinct variations within the binding region. Despite their overall similarity, notable differences are observed between the interaction processes of αCT-M2PI and αCT-nM2PI. Specifically, peaks at N4-Y5, F10-P13, H21, Y42, P50, and R55 are only to the αCT-M2PI complex (Fig. 5E), while peaks at H7, V12, R14, G24, L27, C35, and V58-E59 are present at the αCT-nM2PI interaction alone (Fig. 5F). Shared hotspots are identified at residues L22, L31, F40, C61, L71, and C75 in both mini-proinsulin complexes, suggesting mini-proinsulin variants engage in allosteric behavior to provide a distinct kinetic stabilization strategy with IR, likely driven by the flexible mini-C peptide. Additionally, cross-correlation analysis was conducted to investigate the collective motion between the αCT domain of IR and the mini-proinsulin analogs.

We also performed AlphaFold prediction using nM2PI-IR co-complex sequence to compare if there are distinct conformational alterations of the structure. Minor conformational changes were observed between predicted and docked co-complexes (Fig. S3). Additionally, M2PI and αCT domain shows low predicted local distance difference (pLDDT), offering dynamic regions of the insulin and binding domain, which is in line with our GNM analysis (Fig. 3 and Fig. 5). Interestingly, the αCT domain exhibits strong correlations with the mini-proinsulin variants (Fig. 5B-C), closely similar to the native αCT-INS complex (Fig. 5A), suggesting that despite the mini-proinsulin analogs adopting receptor interaction strategies distinct from that of native insulin, they still manage to obtain favorable and identical structural stability (Fig. 6). Regarding the inspection of the lowest frequency collective motions between the αCT domain of IR and the C-peptides of mini-proinsulin analogs alone, coherent features intrinsically endowed by the rigid blocks within this complex are observed, especially between the aCT and the C-peptide of nM2PI more pronounced (Fig. 6B). This allosteric regulation, again, due to probably the mobility of the C-peptides, might be highlighting the receptor interaction ability of mini-proinsulin analogs to stabilize effectively, even when their methods of achieving such stability diverge from the native form.

**Figure 6.**
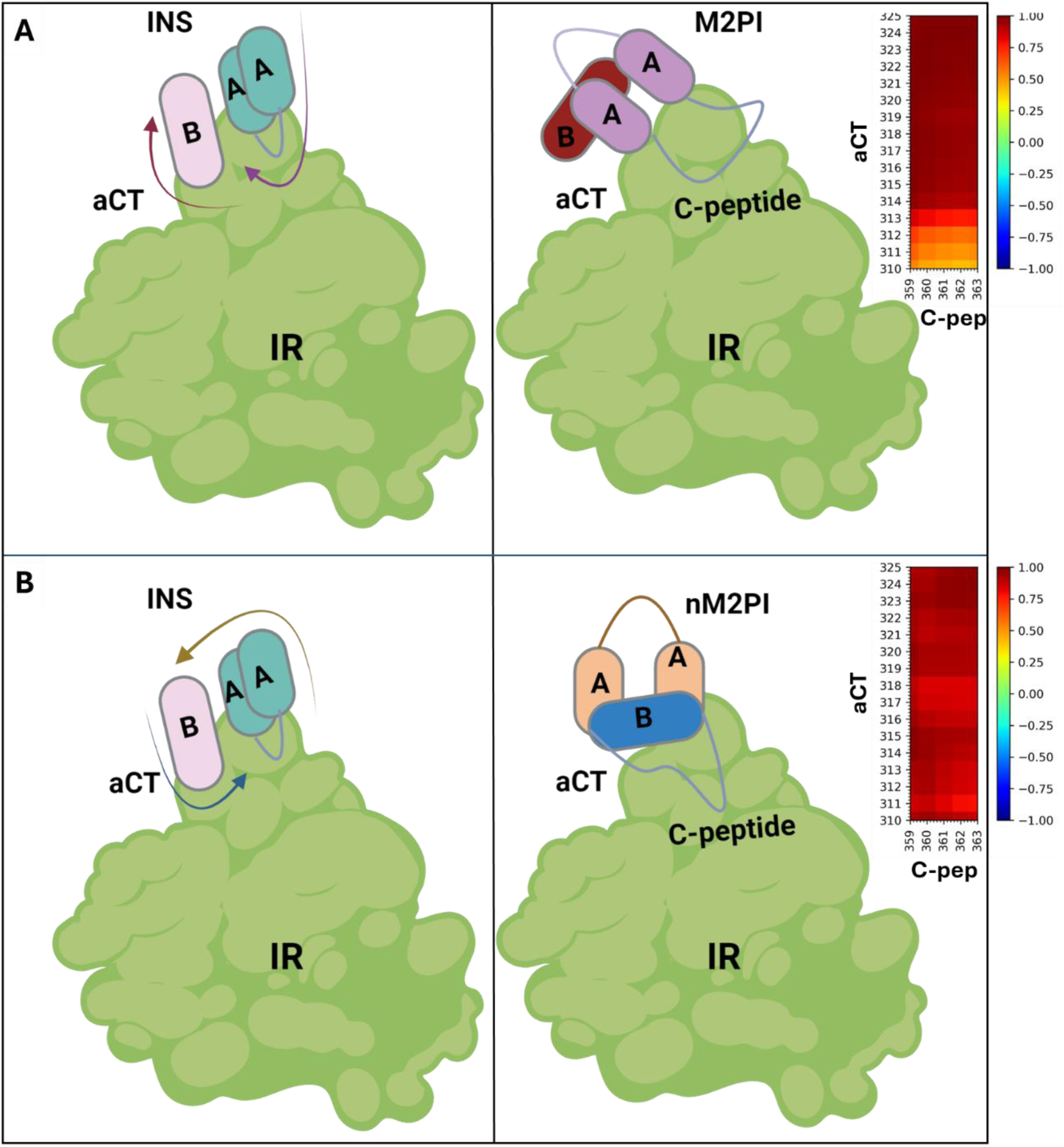
Pictorial representation the allosteric behavior of M2PI and nM2PI compared INS in receptor binding driven by C-peptide mobility. (A) Comparison of how the M2PI position alters its conformation after binding to the αCT receptor domain compared to the INS position within the IR. Cross-correlation analysis by GNM between aCT and C-peptide within the IR-M2PI co-complex. N-terminal of aCT shows at least 50% correlation with C-peptide of IR-M2PI co-complex. (B) Comparison of how the nM2PI position alters its conformation after binding to the αCT receptor domain compared to the INS position within the IR. Cross-correlation analysis by GNM between aCT and C-peptide within the IR-nM2PI co-complex. aCT shows strong correlation with C-peptide of IR-nM2PI co-complex. Residue pairs exhibiting correlated motion, moving in the same direction, are depicted in red regions (C_ijorient_ > 0), while those exhibiting anticorrelated motion, moving in opposite directions, are represented in blue regions (C_ijorient_ < 0). Uncorrelated residue pairs are shown in green (C_ijorient_ = 0.0), as indicated by the color bar in right panels.

## 4. Discussion

The present study computationally models the dynamic interactions between mini-proinsulin analogs and the insulin receptor. Two key rationales underpin the cleaving of the C-peptide in conventional proinsulin. First, optimize the early and late stages of protein refolding pathways [21]. Second, predictions indicate that most dibasic processing sites are located in a β-turn adjacent to regions of α-helix or β-sheet [22]. Hence, introducing a β-turn between these dibasic sites is proposed to facilitate the enzymatic cleavage process required to release the mature insulin, potentially enhancing post-processing efficiency. β-turns are essential structural elements on the protein surface, with their backbone conformation predominantly governed by critical interactions. Recently, Glidden et al. (2017) developed a mini-proinsulin analog (SCI) that C domain contains 6 residues instead of pentapeptide, resulting in a 57-residue protein. In this study, the pentapeptide, YPGDV sequence, was selected due to its notable advantages in promoting β-turn formation. Specifically, studies on β-turns have shown that the presence of proline at the 2^nd^ position and glycine at the 3^rd^ position results in the highest β-turn population. Additionally, the inclusion of deprotonated aspartic acid at the 4^th^ position significantly enhances the stability of the β-turn. The YPGDV sequence has been demonstrated to exhibit a high frequency of β-turn conformations in isolation [11]. Experimental validations reveal that mini-proinsulin incorporating the YPGDV pentapeptide, instead hexapeptide, demonstrates a substantially higher refolding yield than the conventional proinsulin isoform [6, 11]. Previous attempts to construct mini-proinsulin using the RRGSKR linker sequence showed reduced purification efficiency, with unresolved confirmation of correct disulfide bond formation [23]. Conversely, mini-proinsulin with the YPGDV sequence exhibited optimal refolding at a pH range of 11–11.5, which is more alkaline than the conditions typically favored for proinsulin refolding [6]. These results suggest that electrostatic repulsion between the dibasic residues, RR and KR, plays a more pronounced role in mini-proinsulin folding kinetics than full-length proinsulin. Additionally, the differential refolding efficiencies between the YPGDV-based mini-proinsulin and other mini-proinsulin variants with distinct C-peptide sequences become increasingly evident at higher protein concentrations, indicating that YPGDV-based design strategy could serve as a robust alternative for scalable recombinant human insulin production.

Here, we utilized two forms of mini-proinsulin that have YPGDV sequences in C-peptide: the 1^st^ is a previously characterized M2PI structure determined by NMR studies (PDB ID: 1EFE), and the 2^nd^ is a novel designer M2PI variant. The novel variant has been calculated by AF and incorporates specific amino acid substitutions, including R22B to K22B to reduce susceptibility to trypsin degradation, P28B to D28B to mitigate potential aggregation in solution, and N60A to G60A to enhance stabilization of nM2PI in acidic environments (Fig 1). To assess the proper binding of mini-proinsulin analogs to the IR, docking studies were performed using the experimental receptor structure (PDB ID: 6VEP). The results indicate that nM2PI displays higher binding affinity to IR relative to the M2PI structure, likely attributed to the positioning of the C-peptide on the αCT domain (Fig. 1). The binding conformations of the mini-proinsulin analogs diverge substantially from each other and the native IR-INS complex. Normal mode analyses show their (M2PI and nM2PI) capacity to activate the receptor, aligning with experimental evidence indicating that mini-proinsulin analogs keep at least 50% of the receptor-binding activity of native insulin [11]. Interestingly, it was reported that the cellular signaling properties of a novel mini-proinsulin analog, SCI, closely resembled those of WT insulin [10]. Notably, SCI revealed negligible mitogenic activity, indicating a minimal potential to induce uncontrolled cell proliferation. Furthermore, animal studies indicate that SCI demonstrated equivalent potency and duration of action to WT insulin when administered to diabetic rats [10]. Intra-monomer dynamics reveal that M2PI, nM2PI, and SCI exhibit enhanced cooperativity when bound to IR, while increased residual mobility is observed when the analogs are in isolation. Despite these conformational differences, those mini-proinsulin analogs show collective motions that align with those of native INS within the IR, suggesting notable interaction patterns between the mini-proinsulin analogs and IR (Fig. 3 and Fig S2B). These computational characteristics are consistent with experimental findings demonstrating a 20-40% improvement in refolding efficiency [11], and greatly reduced aggregation and thermal propensity [10] compared to native INS.

The global 5 slow ANM modes and local 10 fastest GNM modes reveal that the mini-proinsulin analogs interact with the IR, adopting similar intrinsic dynamics to native INS, yet exhibiting conformational alterations that disrupt conventional interaction pathways and create new ones (Fig. 4 and Fig. 5). Particularly, the local 10 fastest GNM modes identify novel potential hot spots that differ significantly from those observed in the native receptor interaction (Fig. 5), suggesting the mobility of the C-peptide controls the allosteric behavior of mini-proinsulin analogs in receptor binding (Fig 6).

Upon comparing the receptor binding activity of mini-proinsulin analogs with other reported mini-proinsulin variants, it was observed that some designs displayed receptor interaction strengths as low as 0.5% of that of native insulin [24]. For example, insulin cross-linked between A1 and B29 with 2,7-diaminosuberic acid showed substantially reduced receptor binding, similar to proinsulin [25], while a mini-proinsulin with a direct peptide bond between B29 and A1 was effectively inactive [7]. In contrast, the high receptor binding activity exhibited by the experimental mini-proinsulin analogs (M2PI and SCI) in previously available study was supported by our computational analyses. Notably, besides the novel identified potential hot spots, the RR and KR enzyme processing regions were found to play a significant role in receptor interaction. The literature further suggests that these basic residues impact the receptor binding conformation [6]. In M2PI and nM2PI, these residues induce conformational changes within the receptor binding region, potentially contributing to the allosteric modulation of those mini-proinsulins. That is why, in proinsulin, the larger C-peptide likely blocks critical active surfaces on the insulin molecule from interacting with the receptor [26], which may explain the reduced receptor binding affinity observed in proinsulin.

Collectively, despite the ongoing challenges in insulin recombinant production, which remains monopolized by certain companies, and based on the experimental findings, this study suggests that with further experimental validation and optimization, mini-proinsulin analogs hold great potential as viable end products for efficient recombinant insulin production in *E. coli*.

## 5. Conclusion

The primary objective of this study was not to offer nMP12 as a direct alternate for native insulin but to supply computational insights into the structural dynamics and receptor-binding mechanisms of mini-proinsulin analogs. Docking and dynamic analyses aimed to analyze whether the mini-proinsulin analog (nM2PI), despite its structural differences, could perform receptor-binding behavior like native insulin. The computational findings are closely aligned with experimental studies. (i) While a 1998 study reported that mini-proinsulin analogs have approximately 50% of the receptor-binding activity of native insulin, a following study in 2017 showed that a novel designer mini-proinsulin (SCI) revealed potency and duration of action indistinct from WT insulin when administered via intravenous bolus injection in diabetic rats; (ii) SCI also displayed enhanced thermal stability and structural resilience compared to native insulin; and (iii) a 2000 study revealed that a novel designer mini-proinsulin achieved a 20–40% improvement in refolding efficiency compared to proinsulin. The in-silico findings further complement these experimental results by demonstrating (i) Mini-proinsulin analogs exhibit strong receptor-bound dynamics (Fig. 1 and Fig. 2) and collective behavior similar to native insulin in the docked state (Fig. 3 and Fig. 4). (ii) Despite their distinct alternate conformation, mini-proinsulin analogs maintain functional dynamics essential for receptor activation, creating new paths (Fig. 5). (iii) The observed allosteric behavior in mini-proinsulin analogs may contribute to their unique binding mechanisms, offering opportunities for further optimization (Fig. 6).

Collectively, these computational analyses show that nMP12 replicates the biological activity of native insulin, which is also consistent with the experimental studies. It’s also enhanced refolding efficacy, stability under acidic conditions, and potential for cost-effective production in E. coli, highlighting its value in addressing some of the limitations of native insulin, such as stability and manufacturing challenges.

## Data availability

The datasets used and/or analysed during the current study available from the corresponding author on reasonable request.

## Author Contributions

E.A. wrote the entire main manuscript text performing all experiments, computational analyses and preparing all figures.

---

**Supplementary Figure 1.**
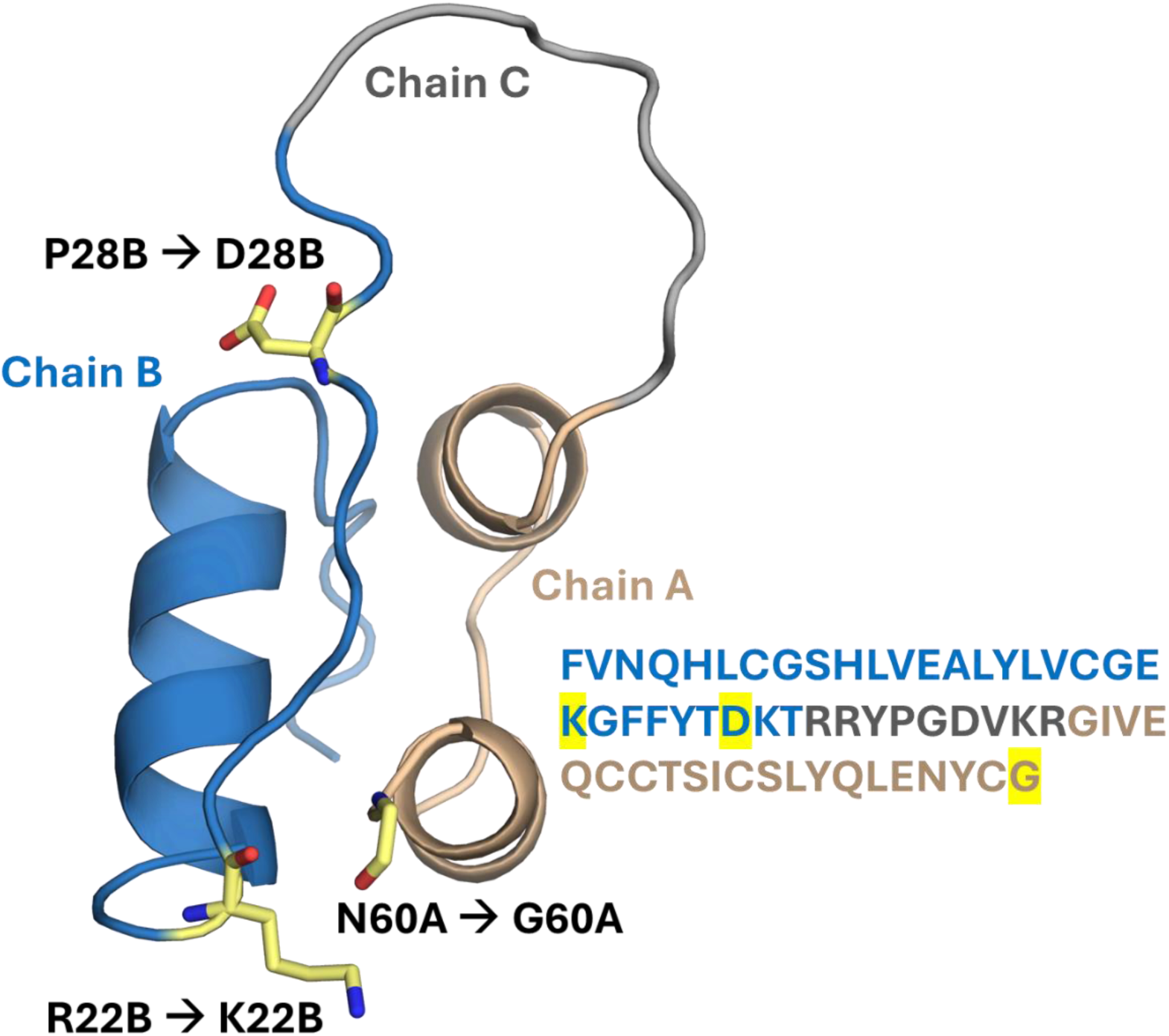
Structural detail of AF predicted mini-proinsulin analog (nM2PI). Chain B is colored in skyblue. Chain A is colored in Chain A. Chain C is colored in gray. Substituted residues are colored in yellow sticks. Proline and Arginine in Chain B have been substituted to Aspartic acid and Lysine. Asparagine in Chain A has been substituted to Glycine.

**Supplementary Figure 2:**
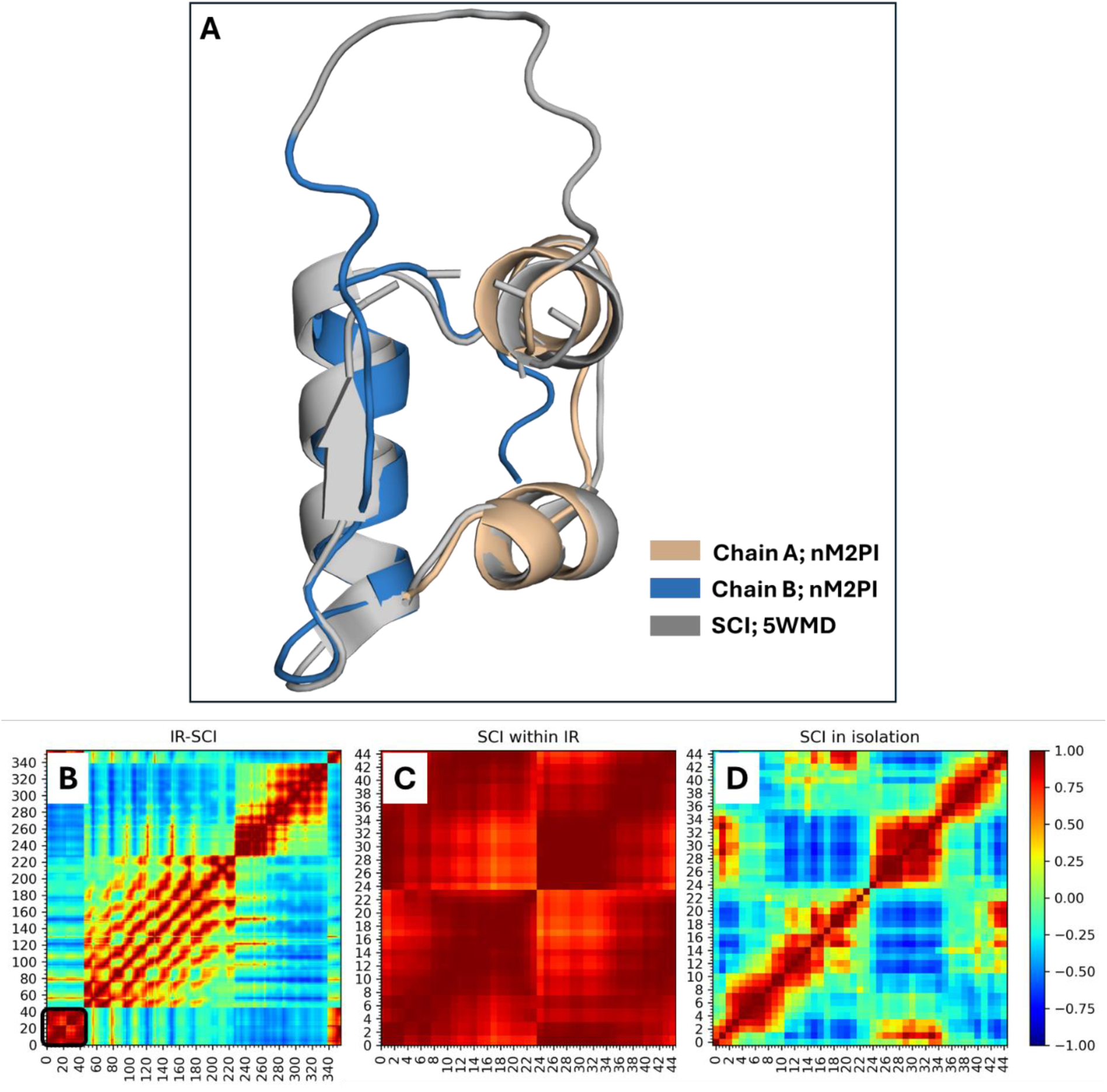
Structural and dynamic comparisons of SCI and nM2PI analogs. (A) Structural overlay of SCI and nM2PI analogs (RMSD of 0.91 Å). (B) Cross-correlation analysis of insulin receptor and mini-proinsulin docking co-complex (referred to as IR-SCI). SCI has been highlighted by black square. (C) Close view of SCI from panel B representing receptor bound mini-proinsulin. (D) Representing the SCI in isolation (unbound to IR).

**Supplementary Figure 3:**
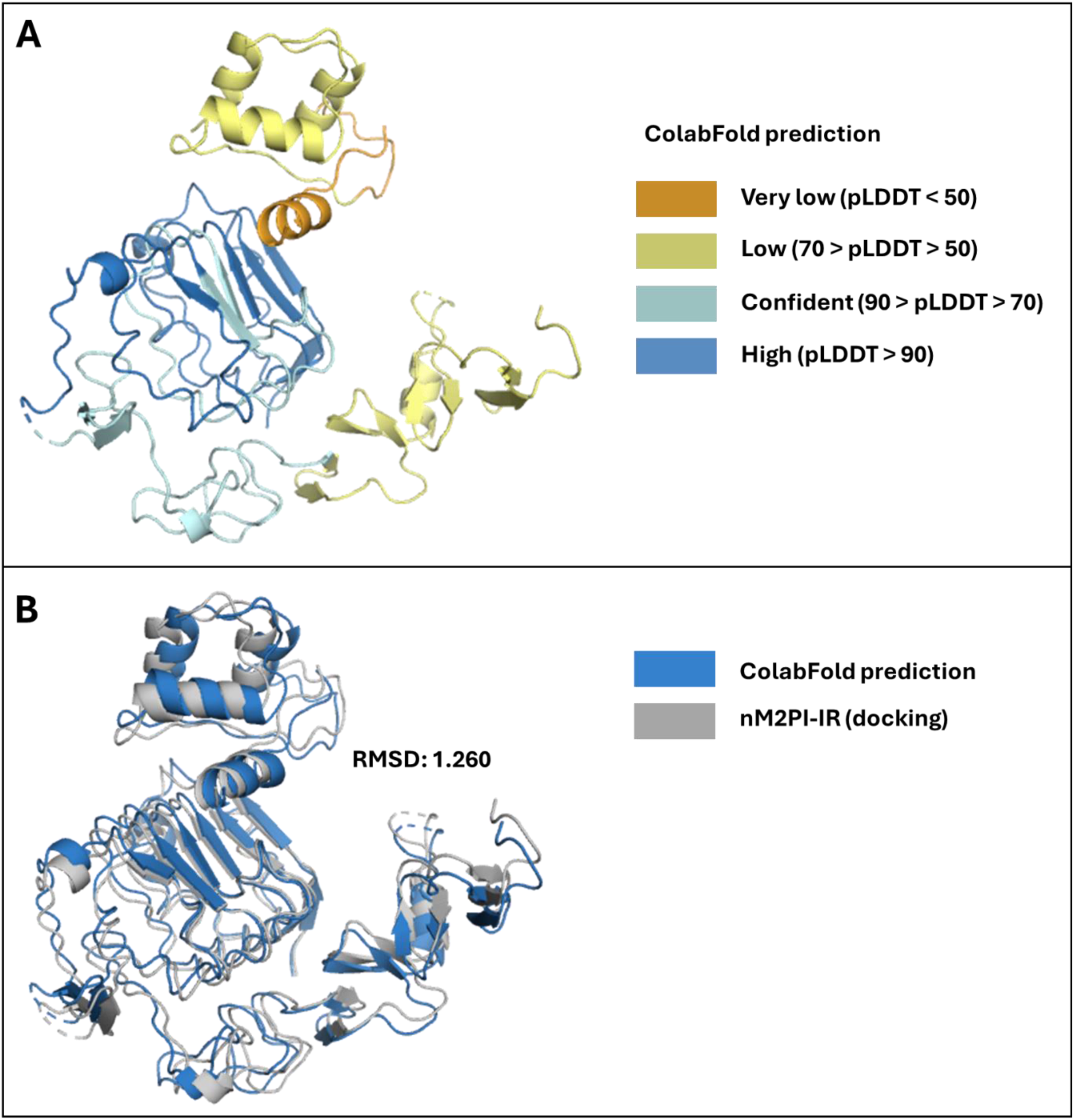
ColabFold prediction of nM2PI-IR co-complex, comparing docked nM2PI-IR co-complex. **(A)** Prediction of nM2PI-IR co-complex is colored in pLDDT (Predicted Local Distance Difference Test) score. nM2PI analog and binding region shows low score, which is in line with our GNM analysis. **(B)** Superposing of predicted vs docked co-complexes shows low RMSD value of 1.260.

